# Mannose receptor (MRC1) mediates uptake of dextran in macrophages via receptor-mediated endocytosis

**DOI:** 10.1101/2024.08.13.607841

**Authors:** Jared Wollman, Kevin Wanniarachchi, Bijaya Pradhan, Lu Huang, Jason G Kerkvliet, Adam D Hoppe, Natalie W Thiex

**Affiliations:** Biology and Microbiology Department, South Dakota State University, Brookings, SD, USA; Chemistry, Biochemistry and Physics Department, South Dakota State University, Brookings, SD, USA

**Keywords:** Mrc1, CD206, mannose receptor, endocytosis, macropinocytosis, macrophage, dextran

## Abstract

Macrophages maintain surveillance of their environment using receptor-mediated endocytosis and pinocytosis. Receptor-mediated endocytosis allows macrophages to recognize and internalize specific ligands whereas macropinocytosis non-selectively internalizes extracellular fluids and solutes. Here, CRISPR/Cas9 whole-genome screens were used to identify genes regulating constitutive and growth factor-stimulated dextran uptake in murine bone-marrow derived macrophages (BMDM). The endocytic mannose receptor c-type 1 (*Mrc1*, also known as CD206) was a top hit in the screen. Targeted gene disruptions of *Mrc1* reduced dextran uptake but had little effect on uptake of Lucifer yellow, a fluid-phase marker. Other screen hits also differentially affected the uptake of dextran and Lucifer yellow, indicating the solutes are internalized by different mechanisms. We further deduced that BMDMs take up dextran via MRC1-mediated endocytosis by showing that competition with mannan, a ligand of MRC1, as well as treatment with Dyngo-4a, a dynamin inhibitor, reduced dextran uptake. Finally, we observed that IL4-treated BMDM internalize more dextran than untreated BMDM by upregulating MRC1 expression. These results demonstrate that dextran is not an effective marker for the bulk uptake of fluids and solutes by macropinocytosis since it is internalized by both macropinocytosis and receptor-mediated endocytosis in cells expressing MRC1. This report identifies numerous genes that regulate dextran internalization in primary murine macrophages and predicts cellular pathways and processes regulating MRC1. This work lays the groundwork for identifying specific genes and regulatory networks that regulate MRC1 expression and MRC1-mediated endocytosis in macrophages.

**Significance Statement:**

- Macrophages constantly survey and clear tissues by specifically and non-specifically internalizing debris and solutes. However, the molecular mechanisms and modes of regulation of these endocytic and macropinocytic processes are not well understood. Here, CRISPR/Cas9 whole genome screens were used to identify genes regulating uptake of dextran, a sugar polymer that is frequently used as a marker macropinocytosis, and compared with Lucifer yellow, a fluorescent dye with no known receptors.
- The authors identified the mannose receptor as well as other proteins regulating expression of the mannose receptor as top hits in the screen. Targeted disruption of *Mrc1*, the gene that encodes mannose receptor, greatly diminished dextran uptake but had no effect on cellular uptake of Lucifer yellow. Furthermore, exposure to the cytokine IL4 upregulated mannose receptor expression on the cell surface and increased uptake of dextran with little effect on Lucifer yellow uptake. Studies seeking to understand regulation of macropinocytosis in macrophages will be confounded by the use of dextran as a fluid-phase marker.
- MRC1 is a marker of alternatively activated/anti-inflammatory macrophages and is a potential target for delivery of therapeutics to macrophages. This work provides the basis for mechanistic underpinning of how MRC1 contributes to the receptor-mediated uptake of carbohydrates and glycoproteins from the tissue milieu and distinguishes genes regulating receptor-mediated endocytosis from those regulating the bona fide fluid-phase uptake of fluids and solutes by macropinocytosis.

## Introduction

Macrophages maintain tissue homeostasis by surveying the tissue fluid space. Using their copious endocytic capacity, macrophages detect and internalize a wide variety of microbial or endogenous molecules via receptor-mediated uptake mechanisms (Kawai and Akira, 2010; Canton *et al*., 2013). In addition, macrophages non-selectively engulf large volumes of fluids and extracellular solutes via macropinocytosis, a bulk fluid-phase uptake mechanism dependent on large, actin-based membrane protrusions (Racoosin and Swanson, 1992; Araki *et al*., 1996; Buckley *et al*., 2020; Quinn *et al*., 2021).

Macropinocytosis has been studied in a variety of cell types and in vivo using fluorescently labelled dextrans as markers of non-specific solute uptake. In macrophages, colony stimulating factor-1 (CSF1) stimulates membrane ruffling and macropinosome formation. CSF1 receptor (CSF1R) ligation by CSF1 promotes phosphatidylinositol 3-kinase (PI3K) activity leading to the formation of phosphatidylinositol (3, 4,5) trisphosphate (PIP3), which can help organize macropinosome formation (Racoosin and Swanson, 1989; Welliver and Swanson, 2012; Maekawa *et al*., 2014). Macropinosomes are important not only for internalizing extracellular solutes including antigens, pathogens, and nutrients, but also for directing receptor traffic within the cell. For example, CSF1R is degraded following trafficking to macropinosomes while integrins are recycled from macropinosomes (Lou *et al*., 2014; Freeman *et al*., 2020).

The discovery of the bacterial CRISPR/Cas9 genome-editing system and its adaptation to mammalian cells has facilitated precise genome editing and facilitated whole-genome screening to discover novel regulators of a variety of eukaryotic cellular processes (Jinek *et al*., 2012; Cong *et al*., 2013; Hsu *et al*., 2013; Lander, 2016). Pooled sgRNA libraries targeting the majority of genes within a genome enable whole-genome forward genetic approaches allowing users to efficiently identify genes regulating specific cellular activities or phenotypes (Shalem *et al*., 2014; Haney *et al*., 2018).

Here, we developed a slate of CRISPR/Cas9 whole-genome screens to identify genes regulating dextran internalization in bone marrow-derived macrophages (BMDM). We used the Brie pooled sgRNA library to disrupt gene function (Doench *et al*., 2016). Cells were sorted based on their ability to internalize fluorescent dextran in the presence or absence of CSF1 stimulation and identified sgRNAs that enhanced or reduced the ability of cells to internalize dextran. A total of 688 genes were identified as regulators of dextran uptake in BMDMs with false discovery rates (FDR) less than 0.1. The endocytic receptor macrophage mannose receptor (MRC1 or CD206) was a significant hit in both unstimulated and CSF1-stimulated dextran uptake. In addition, genes regulating MRC1 expression and trafficking were observed to be critical for regulating dextran internalization. These results demonstrate that macrophages internalize dextran via a receptor-mediated mechanism, which is a significant finding for the field of macropinocytosis as dextrans have been assumed to be internalized via non-specific mechanisms and have been extensively used to study macropinocytosis.

## Results

### CRISPR/Cas9 whole-genome screen identifies Mrc1 as a positive regulator of fluorescent dextran uptake in BMDM

CRISPR/Cas9 whole-genome screens were implemented to identify and rank genes regulating both constitutive and CSF1-stimulated fluorescent dextran uptake (Figure 1). BMDMs transduced with the Brie sgRNA library were exposed to fluorescent dextran and the highest and lowest 20% of cells were sorted by flow cytometry (Joung *et al*., 2017). The sgRNA inserts in the sorted cell populations were then amplified by PCR and deep sequenced providing read counts for each sgRNA in the library in the low- and high-fluorescence cell populations. MAGeCK was used to calculate log2-fold change (LFC) between the low-fluorescence and high-fluorescence populations by gene (Li *et al*., 2014, 2015b). In unstimulated BMDMs, the screens identified 561 genes with a false discovery rate (FDR) less than 0.1 that regulate constitutive dextran uptake in the absence of CSF1 (Figure 2A, Supplemental Table S1). The negative LFC indicate genes where the sgRNA was higher in the low-fluorescence population than the high-fluorescence population identifying genes that positively regulate dextran uptake. The genes with the most negative LFC were *Atp6v0a1*, *Slc35a2*, *Ptpn6*, *Mrc1* and *VPS33a. Phf5a*, *Nprl3*, *Syk, Vmn1r58* and *Stk11* were the genes with the largest positive LFC, indicating gene disruptions that increased dextran uptake suggesting that the gene product provides negative regulation of dextran uptake (Figure 2A, Supplemental Table S1). Since growth factors increase bulk fluid and solute uptake in macrophages by increasing macropinocytic activity (Racoosin and Swanson, 1989; Pacitto *et al*., 2017), we also identified genes that positively or negatively regulate dextran uptake in response to CSF1 stimulation. A total of 353 genes with an FDR less than 0.1 were identified (Supplemental Table S2). The positive regulator genes with the most negative LFC (low fluorescence) were *Mrc1, Ptpn6, Slc35a2, Mtmr6,* and *Atp6v1e1.* Whereas the negative regulators of CSF1-stimulated dextran uptake were *Vmn1r149, Trem2, Tyrobp, Nprl2,* and *Tsc1* (Figure 2B, Supplemental Table S2). Other top hits of interest are indicated on the volcano plots (Figures 2A and B).

**Figure 1.**
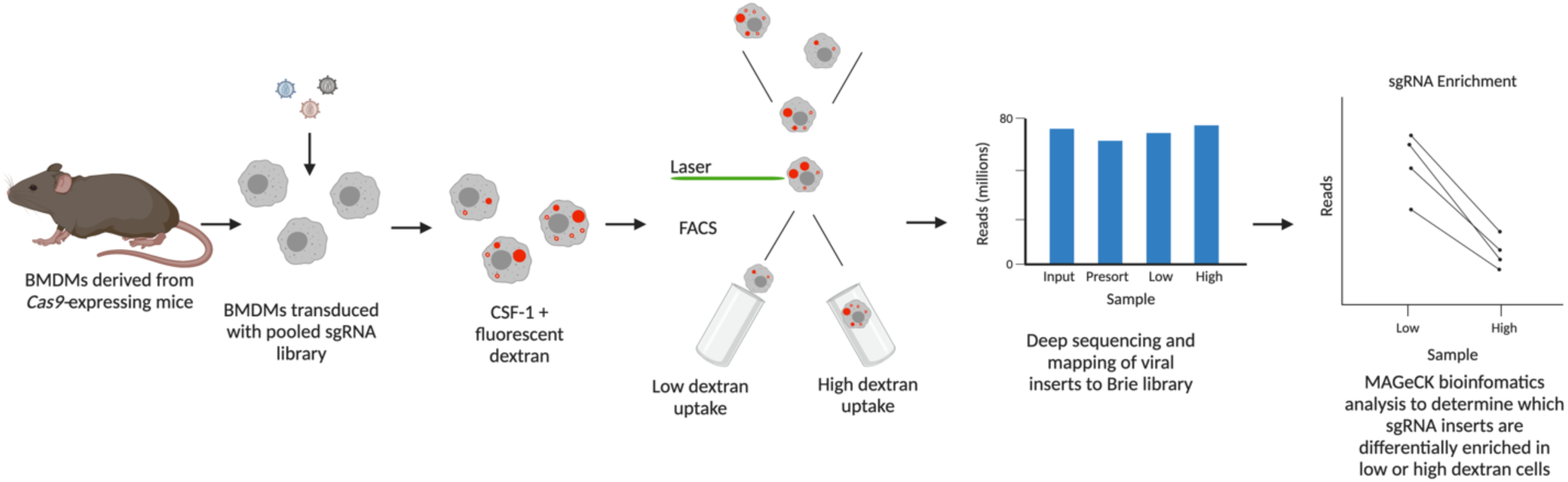
CRISPR/Cas9 screen workflow to identify genes regulating dextran uptake in bone marrow- derived macrophages. BMDMs were derived from Cas9-expressing mice, transduced with the Brie sgRNA library, selected with antibiotic, and cultured for 10 days to allow time for gene disruptions and protein turnover. The day before sorting, cells were starved of CSF1 overnight. Cells were exposed to 40 kDa fluorescent dextran in the presence or absence of CSF1 for 15 min. The upper and lower quintiles of cells were sorted by fluorescence, genomic DNA was purified from low- and high-drinker populations, and bar-coded sgRNA inserts were amplified and sequenced. The sgRNA insert sequences were mapped to the sgRNA library, and statistical enrichment of sgRNA inserts in the low- and high-fluorescence groups was determined using the MAGeCK bioinformatics tool. Created with BioRender.com.

**Figure 2.**
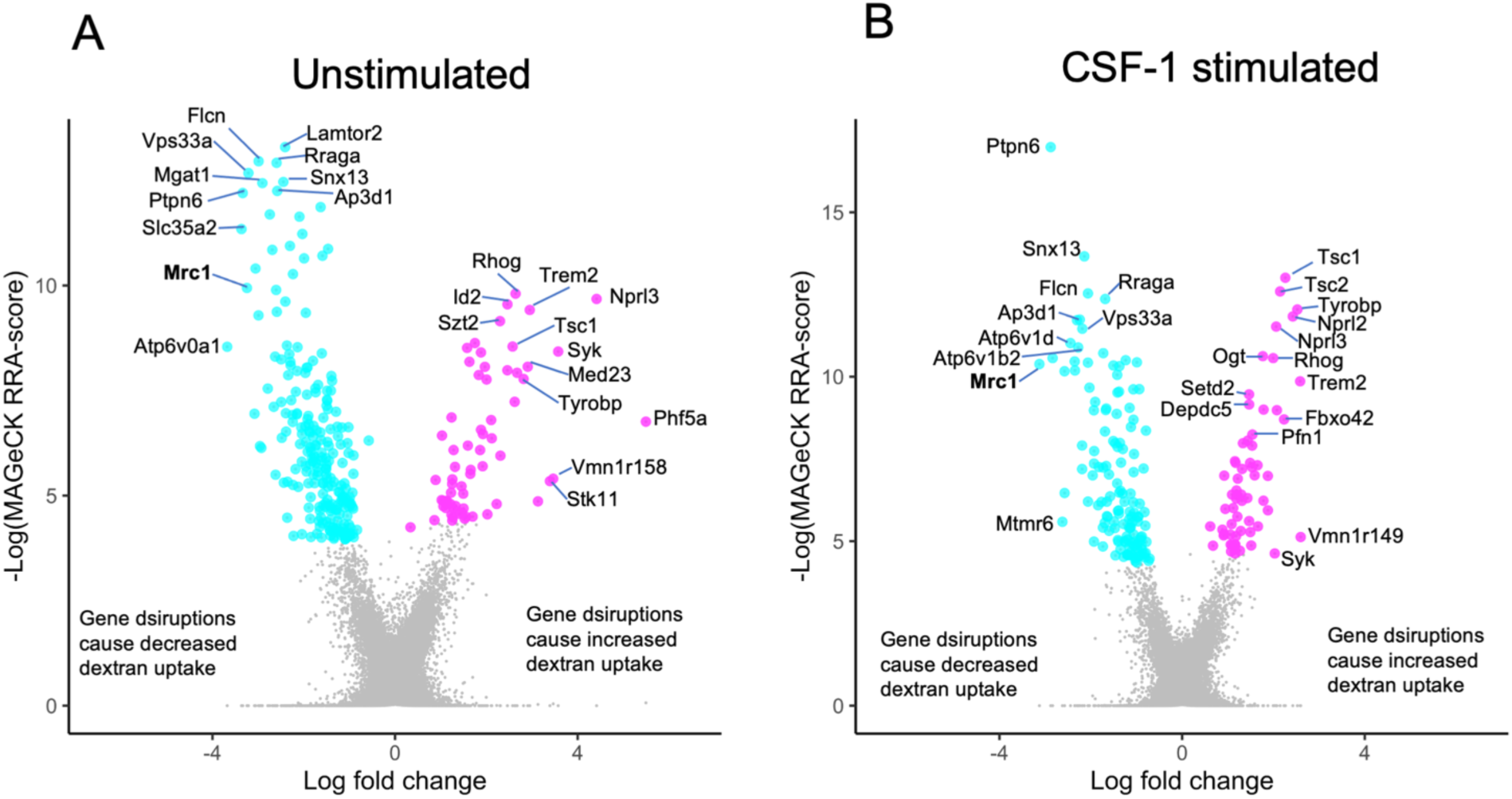
Mrc1 predicted as a regulator of dextran uptake in BMDM by CRISPR/Cas9 whole genome screens. Volcano plots show the MAGeCK RRA score plotted against the log2-fold change of sgRNA enrichment in the high- vs low-dextran uptake populations for each gene. Genes where Cas9-mediated disruptions diminish dextran uptake are shown in cyan and those that increase dextran uptake in magenta (FDR < 0.1). A) Populations of BMDM transduced with the BRIE sgRNA library were exposed to dextran for 30 min and sorted into high and low populations in the absence of CSF1 to identify genes regulating constitutive regulation of dextran uptake. B) BMDM transduced with the BRIE sgRNA library were exposed to dextran in the presence of CSF1 for 30 min and then sorted into high and low fluorescence populations to identify genes regulating growth factor stimulated macropinocytosis.

### The endocytic receptor MRC1 confirmed as a key mediator of dextran uptake in BMDMs

The Brie sgRNA library contains multiple sgRNA for each gene. We selected genes of interest and plotted normalized read counts from the next generation sequencing for each sgRNA in the low- and high-fluorescence populations. Comparison of sgRNA read counts between the cells sorted into the low- dextran and high-dextran populations showed high consistency among the different guides from the same gene (Figure 3A) providing confidence in the hit calling. BMDM containing *Mrc1, Vps33a* or *Ptpn6* sgRNA inserts were more likely to sort into the low-dextran fluorescence populations suggesting that these genes encode proteins that are positive regulators of dextran uptake. In contrast, BMDM containing *Syk* or *Rhog* sgRNA were more likely to sort into the high-dextran uptake population suggesting these genes are inhibitors or negative regulators of dextran uptake.

**Figure 3.**
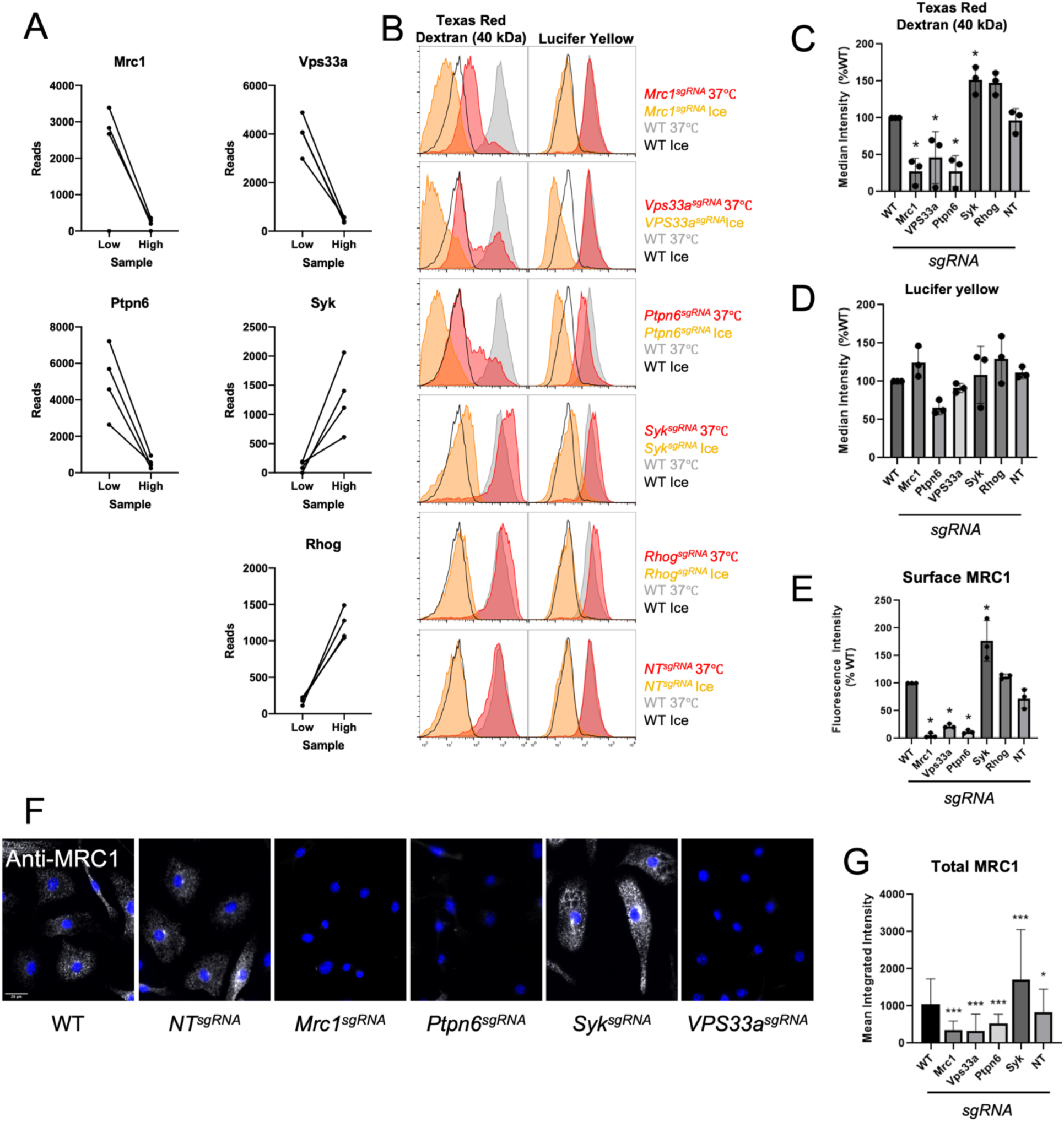
Dextran uptake and Lucifer yellow uptake are differentially regulated in BMDM. A) The normalized read counts from the CSF1-stimulated screen of individual sgRNAs within the low- and high- dextran sorted cells for select genes. B) BMDMs with the indicated targeted gene disruptions were starved of CSF1 overnight and then stimulated with CSF1 for 30 min in the presence of both Lucifer yellow and 40 kDa Texas red dextran at 37°C or on ice. Each histogram compares WT BMDM (gray/black) with the indicated targeted gene disruption (red/orange). NT = non-targeting sgRNA C) Replicates of dextran uptake by genotype. Dots are median fluorescence intensity of dextran uptake at 37°C sample minus the median of a corresponding no-dye control to account for differences in autofluorescence. Bar graph shows the mean of 3 replicate experiments, error bars are SD. * p < 0.05. D) Replicates of Lucifer yellow uptake by genotype. Dots are median of Lucifer yellow uptake at 37°C sample minus the median of the genotype-matched no dye sample. Bar graph shows the mean of 3 replicate experiments, error bars are SD. E) BMDM with the indicated gene disruptions were immunostained for surface MRC1 and fluorescence intensity analyzed by flow cytometry. Bar graph shows the mean of 3 replicate experiments; error bars are SD; * p<0.05. F) BMDM with the indicated gene disruptions were fixed, permeabilized and stained for total MRC1 (white) and nuclear stain (blue). Scale bars show 20 µm. NT = non-targeting sgRNA. G) Quantification of F. * p< 0.05

Dextran is frequently used as a marker of bulk, non-specific uptake to study macropinocytosis in macrophages and other cell types (Racoosin and Swanson, 1992; Redka *et al*., 2018; Lin *et al*., 2020; Le and Machesky, 2022). However, screen analyses predicted the endocytic receptor MRC1 as a strong positive regulator of both constitutive and CSF1-stimulated dextran uptake suggesting that macropinocytosis may not be the main route of dextran uptake in BMDM. MRC1 is a C-type lectin that binds terminal sugars such as mannose on proteoglycans of both endogenous and exogenous origin and internalizes them via a clathrin-dependent mechanism (Gazi and Martinez-Pomares, 2009), and dendritic cells internalize dextrans via MRC1-mediated mechanisms (Sallusto *et al*., 1995; Hackstein *et al*., 2002). To validate the genes identified by the screen and to determine whether BMDMs take up dextran by a receptor-mediated mechanism or a non-specifically by macropinocytosis, we made CRISPR- Cas9 targeted gene disruptions in *Mrc1* as well as other genes highly ranked by the screen *(Vps33a, Ptpn6, Syk,* and *Rhog)*. Based on performance in the screen we selected sgRNAs from the Brie library to make targeted gene disruptions for further validation (Supplemental Table S3). We observed that transducing Cas9-expressing BMDM with sgRNAs targeting these genes recapitulated the low- or high- dextran uptake phenotype predicted by the screens, (Figure 3B and C). Specifically, *Mrc1^sgRNA^*, *Vps33a^sgRNA^, and Ptpn6^sgRNA^* BMDM displayed a loss of the ability to efficiently internalize Texas Red dextran (40 kDa) at 37°C, whereas *Syk ^sgRNA^* targeted cells displayed an enhanced ability to take up dextran (Figure 3B and C), recapitulating the phenotype predicted by the screen. This observation was further supported using a temperature block (incubation on ice) to prevent both endocytosis and macropinocytosis while allowing binding to external receptors. *Mrc1 ^sgRNA^*, *Vps33a ^sgRNA^* and *Ptpn6 ^sgRNA^* BMDMs bound less dextran than WT BMDM (Figure 3B).

Given that *Mrc1* gene disruption had such a profound effect on dextran binding and uptake and that this receptor is known to bind glycans, we compared the uptake of Lucifer yellow across these same mutants by measuring the simultaneous uptake of Texas red dextran and Lucifer yellow (left vs. right columns of Figure 3B). Lucifer yellow has a low molecular weight (442 g/mol), negative charge (-2e), is polar, and has been demonstrated to be a bona fide fluid-phase marker (Swanson *et al*., 1985). Indeed, unlike dextran uptake, which is decreased in *Mrc1^sgRNA^*and *Vps33a^sgRNA^* BMDM, no difference in Lucifer yellow uptake was observed in these mutants compared with *non-target^sgRNA^*(*NT^sgRNA^*) and wildtype BMDMs. (Figure 3D). These data suggest that Lucifer yellow and dextran, which were mixed together before addition to the cells, were internalized by different cellular processes and that MRC1 mediates dextran but not Lucifer yellow uptake (Figure 3B-D).

To determine if changes in MRC1 expression contribute to the defects observed in dextran uptake in these mutants, we quantified its cell surface expression by flow cytometry (Figure 3E) and total expression by immunofluorescence microscopy (Figure 3 and G). Indeed, *Mrc1^sgRNA^* cells were devoid of the receptor (Figure 3F and G) as were *Vps33a* ^sgRNA^ and *Ptpn6^sgRNA^* BMDMs (Figure 3F and 3G) consistent with lack of MRC1 expression leading to decreased dextran binding and internalization.

### Receptor-mediated endocytosis is the primary mode of dextran uptake in BMDM

Previous reports indicated that the size of the dextran may impact the route of internalization with 70 kDa dextrans being internalized via a clathrin- and dynamin-independent mechanism in HeLa cells and coelomocytes (Li *et al*., 2015a). Therefore, to determine if dextran size influenced its uptake in a MRC1- dependent manner, we compared uptake of 3, 40 or 70 kDa Texas red dextrans in WT and *Mrc1^sgRNA^* BMDM (Figure 4A-D). CRISPR-based targeted disruptions of *Mrc1* in BMDM and protein depletion were confirmed by immunoblot (Supplemental Figure S1). Uptake (37°C) and binding (ice) of all sizes of dextran was strongly influenced by MRC1, whereas Lucifer yellow was largely unaffected (Figure 4A). A weak size dependence was observed for the uptake of dextran in *Mrc1^sgRNA^* BMDM at 37°C where *Mrc1^sgRNA^* gene disruption had a bigger effect on 70 kDa dextran uptake than 3 kDa uptake. This finding is the opposite of what would be expected if 70 kDa dextrans were size excluded from endosomes and demonstrates that 70 kDa dextrans are efficiently taken up by MRC1-mediated mechanisms. Treatment with Dyngo4a dramatically reduced dextran uptake indicating a critical role for dynamin on its internalization but had little effect on Lucifer yellow (Figure 4E). The clathrin inhibitor, Pitstop2, strongly disrupted the uptake of both dextran and Lucifer yellow (Figure 4E). Taken together, these results are consistent with dextran uptake being mediated by a dynamin-dependent small vesicle endocytosis and further indicates that dextrans and Lucifer yellow are internalized by different mechanisms in macrophages. Whereas CSF1 stimulation of macropinocytosis strongly enhanced the uptake of Lucifer yellow (∼5x), it had only a slight effect on the uptake of dextran (Figure 4E) supporting the assertion that macropinocytosis is a minor route for uptake of dextran in macrophages.

**Figure 4.**
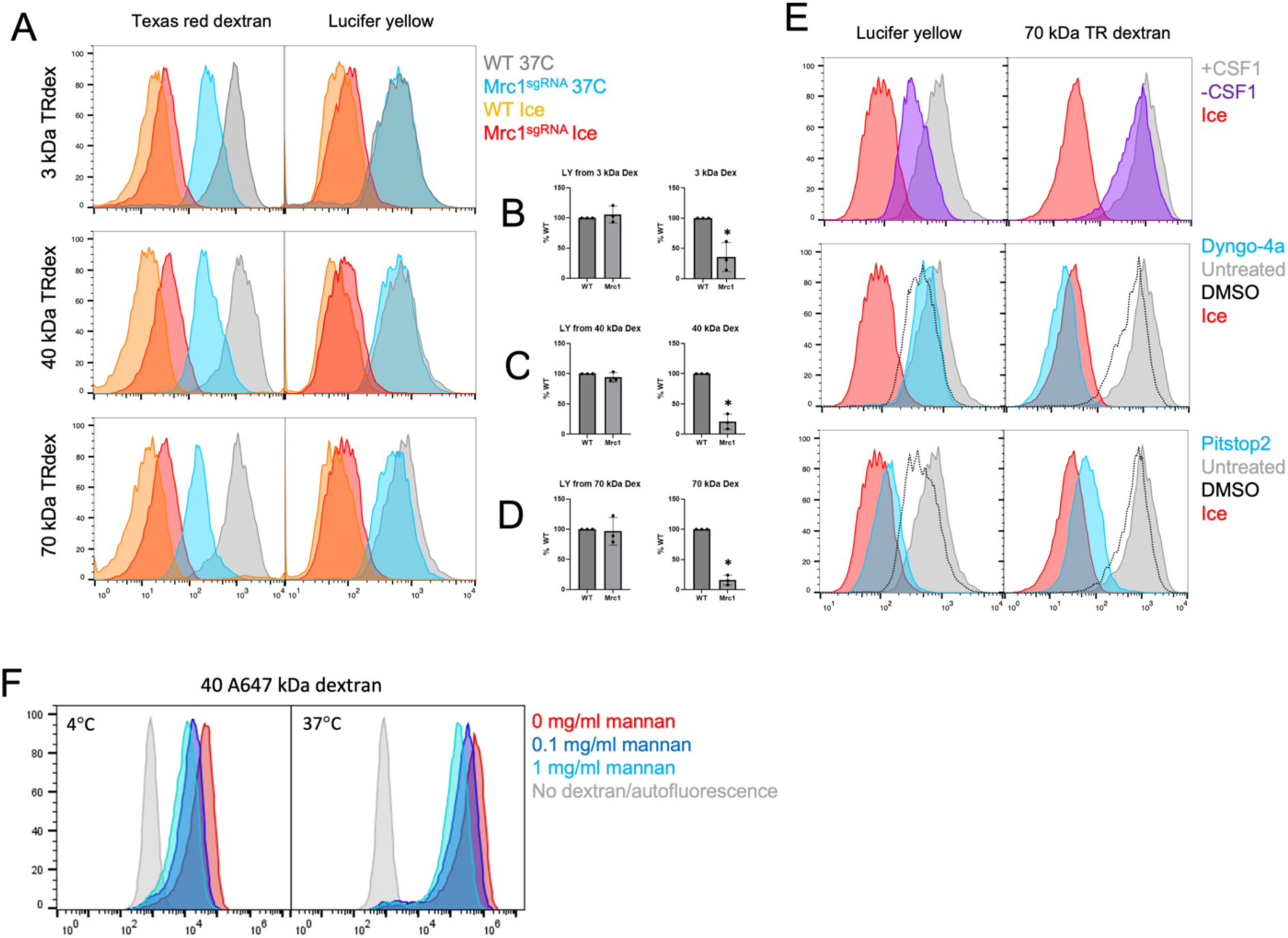
Disruption of endocytosis decreases dextran uptake in BMDM A) Mrc1sgRNA BMDM have impaired uptake of 3, 40 and 70 kDa TR dextran but not Lucifer yellow. BMDM were starved of CSF1 overnight and then exposed to Lucifer yellow and indicated sizes of Texas red dextran (250 ug/ml) at 37°C to measure dye uptake or on ice to measure dye binding. B-D) Plots of medians of three replicate experiments comparing uptake of 3, 40 and 70 kDa dextrans and Lucifer yellow uptake in WT and Mrc1sgRNA BMDM. Error bars are SD; * p< 0.05. E) WT BMDM were starved of CSF1 overnight pretreated with the indicated treatments and then exposed to mixed Lucifer yellow and 70 kDa dextran in the presence of CSF1. F) Mannan causes a dose-dependent inhibition of dextran binding and uptake. BMDM were starved of CSF-1 overnight followed by a 30-min pretreatment with 0, 0.1, or 1 mg/ml mannan. Then, cells were stimulated in the presence of CSF1 for 30 min at 4°C or 37°C in the presence of 0.1 mg/ml AF647 (40 kDa) to measure dextran binding and dextran uptake.

To further test the requirement of MRC1 in dextran uptake, we pre-treated cells with mannan, a polymer of mannose, which binds to MRC1 and has previously been observed to be a competitive inhibitor of MRC1 ligands including dextran, horseradish peroxidase and ovalbumin (Nakajima and Ballou, 1974; Montaner *et al*., 1999; Autenrieth and Autenrieth, 2009). Indeed, pretreatment of BMDMs with increasing concentrations of mannan decreased dextran binding to the cell surface on ice and decreased dextran internalization at 37°C (Figure 4F) indicating competition for receptor binding sites.

### Dextran is internalized by Mrc1 on small endosomes and partially by macropinocytosis

We next sought to reconcile dextran’s historical use as a bulk fluid-phase marker and our finding that MRC1 mediates its uptake. Early studies of macropinocytosis used high concentrations of dextran at about 1 mg/mL, which we speculated may saturate MRC1 with the remainder being internalized through macropinocytosis. Following treatment with mannan, CSF1-stimulated macrophages were allowed to take up Lucifer yellow mixed with 100 ug/ml or 500 ug/ml Texas red 40 kDa dextran (Figure 5A).

**Figure 5.**
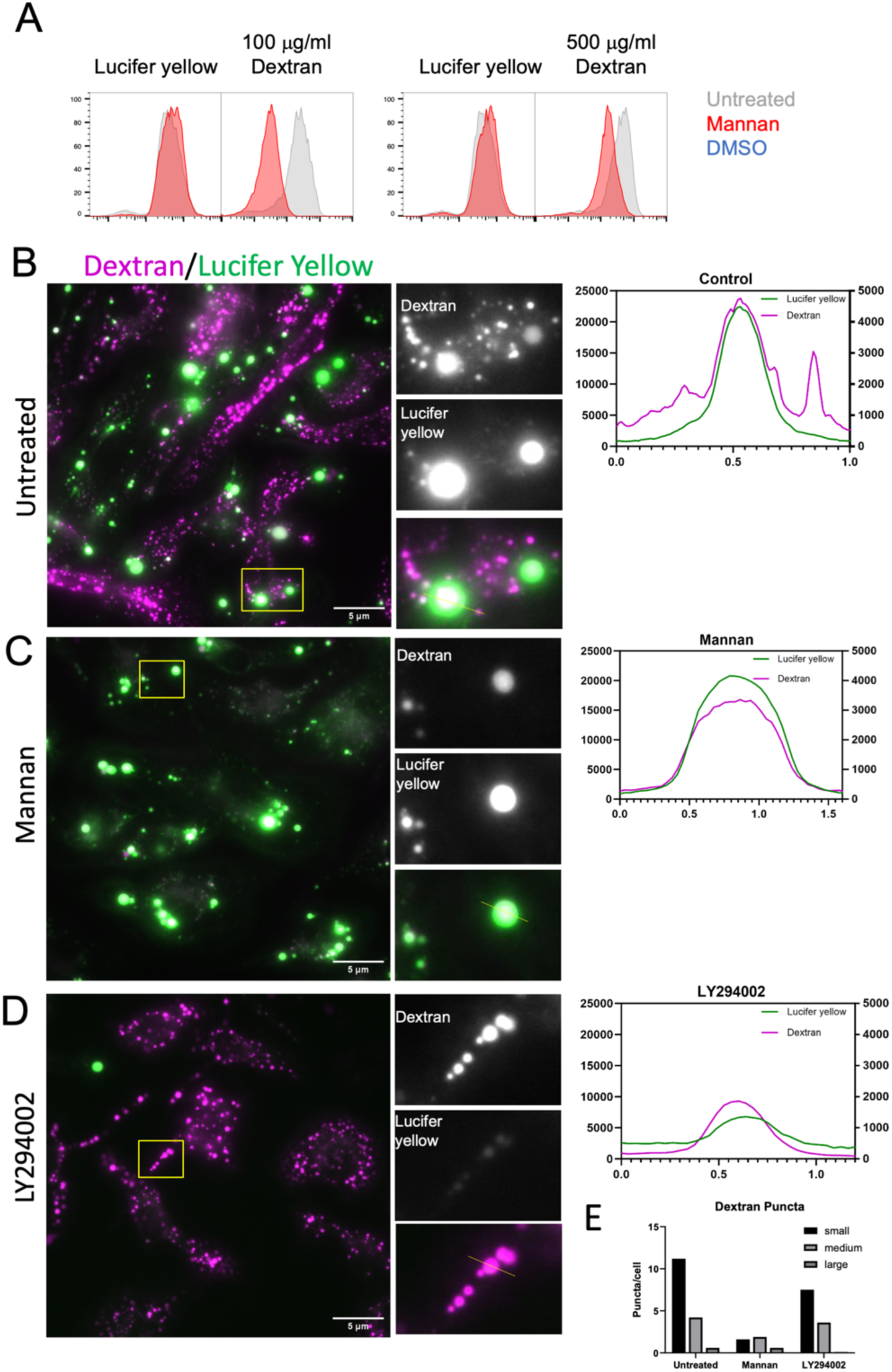
Both macropinocytosis and endocytosis mediate dextran uptake in BMDM. A) BMDM were starved overnight of CSF1 and then pre-treated with either 1 mg/ml mannan, 50 µM LY294002 or DMSO (solvent control for LY294002) for 30 min before stimulation with CSF1 in the presence of both Lucifer yellow and 100 or 500 µg/ml Texas red 40 kDa dextran for 30 min. Following stimulation, cells were washed and analyzed by flow cytometry. B) BMDM were starved overnight of CSF1 and then restimulated with CSF1 in the presence of both Lucifer yellow and 40 kDa Texas Red dextran, washed and imaged. Line scans (yellow lines on insets) of fluorescence intensity to compare relative intensity of dextran and Lucifer yellow in small and large puncta. C) Lucifer yellow and Texas red dextran uptake in the presence 1 mg/ml mannan. D) Lucifer yellow and Texas red dextran uptake in the presence of of with 50 µM LY294002. E) Quantification of the size of dextran puncta/cell with small (< 0.3 um) medium (0.3-1.1 um) or large > 1 um in untreated (n= 49), mannan (n=35) or LY294002 (n=49) treated BMDM.

Pretreatment with mannan significantly reduced dextran uptake at both low and higher concentrations of dextran but did not influence Lucifer yellow uptake (Figure 5A). Specifically, pre-treatment with mannan reduced uptake of 100 µg/ml Texas red dextran by over 90% and reduced uptake of 500 µg/ml Texas red dextran by approximately 60%. Thus, mannan pre-treatment had a greater effect on uptake of lower concentrations of dextran as would be expected with a saturatable uptake mechanism such as receptor-mediated endocytosis as seen by a bigger effect of mannan competitive inhibition at 100 µg/ml dextran compared to 500 µg/ml.

To visualize the vesicles containing dextran and Lucifer yellow following treatment, we observed CSF-1 stimulated Lucifer yellow and fluorescent dextran uptake by epifluorescence microscopy (Figure 5B-D) in untreated, mannan and LY294002-treated BMDM. In the untreated BMDM, three classes of puncta were observed: small (< 0.3 μm), medium 0.3-1.1 μm), and large (> 1.1 μm). Lucifer yellow was observed to be predominantly localized to large puncta (macropinosomes) along with dextran, whereas small puncta were highly enriched dextran suggesting a receptor mediated uptake mechanism rather than bulk non-specific uptake (Figure 5B). Mannan treatment greatly diminished dextran uptake in small macropinosomes (Figure 5C) consistent with lack of receptor-mediated endocytic uptake under that condition. The PI3K inhibitor LY294002 inhibited the formation of large vesicles consistent with the drug causing a defect in macropinocytosis (Figure 5D). Analysis of the sizes of dextran puncta (Figure 5E) illustrated that mannan was highly efficient at preventing dextran uptake into small and medium vesicles, whereas LY294002 prevented dextran uptake in large vesicles. Together, these data suggest mannan inhibits receptor-mediated endocytosis of fluorescent dextran while LY294002 inhibits fluid-phase uptake.

### IL4 enhances dextran uptake in murine macrophages by upregulating MRC1

The anti-inflammatory cytokine IL4 causes alternative activation of macrophages and increases MRC1 expression in human monocyte-derived macrophages. IL4 has been reported to stimulate macropinocytosis (Redka et al., 2018; Montaner et al., 1999). Here we show that IL4 treatment dramatically increased MRC1 expression on the cell surface of BMDMs (Figure 6A), and therefore asked whether increase in dextran uptake in IL4-treated macrophages can be attributed to greater expression of MRC1. We used flow cytometry to measure both Lucifer yellow and dextran uptake in IL4-treated WT and Mrc1^sgRNA^ BMDM following CSF1 stimulation, and whether mannan inhibited dextran uptake in either genotype. IL4-treated cells took up a similar amount of Lucifer yellow as non-IL4-treated cells but took up over four times more dextran than untreated WT cells (Figure 6B and C). The increased dextran uptake in IL4-treated cells was abolished by mannan, suggesting that increased uptake of dextran by IL4- treated BMDM is via increased MRC1 protein and receptor-dependent dextran uptake, rather than macropinocytosis. Furthermore, Mrc1^sgRNA^ BMDM showed no increase in dextran uptake following IL4 stimulation and were unaffected by competition with mannan suggesting the expression of MRC1 is critical to manifest the effects of IL4 on dextran uptake (Figure 6B and C). Using epifluorescence microscopy, we observed dextran uptake in IL4-treated cells following CSF1 stimulation to confirm that IL4-treated cells internalize dextran in small, bright puncta, which are assumed to be endosomes and large, dimmer macropinosomes which contain Lucifer yellow (Figure 6D). Together, these data support the claim that IL4-treated cells increase dextran uptake by upregulating MRC1 expression rather than by appreciable upregulation of macropinocytosis.

**Figure 6.**
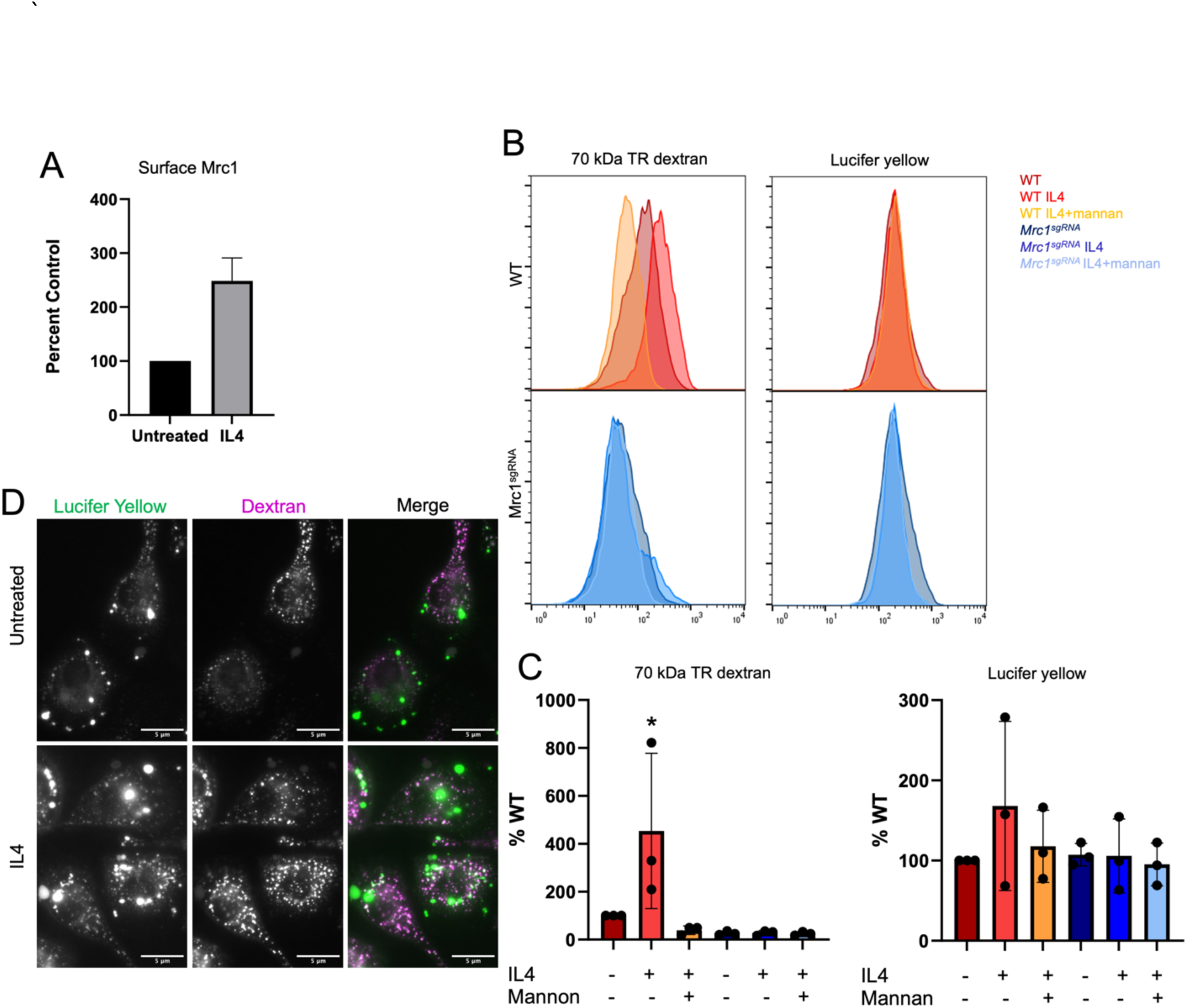
IL4-treated BMDM take up increased levels of fluorescent dextran by upregulating expression of MRC1. A) Surface staining of MRC1 in BMDM were treated with 50 ng/ml IL4 for 48 h and compared to untreated cells. B) Effect of IL4 with or without mannan on dextran uptake in WT and Mrc1sgRNA BMDM. Cells were treated with 50 ng/ml IL4 for 48 h before exposure to Texas red dextran mixed with Lucifer yellow for 30 min C) Plots of medians of three replicate experiments comparing uptake of 70 kDa dextran and Lucifer yellow uptake in WT and Mrc1sgRNA BMDM. Error bars are SD; * p< 0.05. D) Primary bone marrow macrophages were treated with 50 ng/ml IL4 for 48 h and starved overnight of CSF1 using DMEM plus 10% FBS. Cells were then stimulated with CSF1 in the presences of 0.08 mg/ml A647 40 kDa dextran and 0.5 mg/ml Lucifer yellow in HBSS for 5 min and then washed and imaged in HBSS.

## Discussion

Using CRISPR/Cas9 whole-genome screens, we identified MRC1 as a major factor contributing to the uptake of dextran in BMDMs. This work revealed that dextran, which is widely used to study macropinocytosis, has minimal utility as a marker for this process when MRC1 is expressed. Moreover, our work challenges the prevalent notion that high molecular weight dextran internalization is restricted to large compartments, but rather, is readily taken up by small vesicle receptor-mediated endocytosis. This is consistent with the fact that the diameter of 70 kDa dextran is 3.4 nm in size, which easily fits within a 200 nm endosome (Berthiaume *et al*., 1995). Moreover, we demonstrate that Lucifer yellow identifies non-specific uptake by macropinocytosis with high fidelity.

The screens identified genes that may be critical for regulating MRC1 endocytic activities and provide insight into how MRC1 may contribute to surveillance of host and pathogen derived molecules. Specifically, as a pattern recognition receptor, MRC1 can internalize both endogenous and pathogenic molecules containing mannose, fucose, N-acetyl glucosamine, or glucose polymers (Feinberg et al., 2000; Iobsts and Drickamero, 1994). We found that the endocytosis of dextran by MRC1 was mediated by a dynamin and clathrin inhibitable mechanism and competitively inhibited by mannan, a known ligand of MRC1. Moreover, the screen identified *Ap2s1*, a gene encoding a clathrin adaptor protein important for clathrin assembly, as a positive regulator, further implicating clathrin-mediated endocytosis in dextran uptake (Kovtun *et al*., 2020). Thus, host and pathogen associated molecules recognized by MRC1 will likely quickly acidify and merge into the endo-lysosomal system. Notably, our screen identified 14 vacuolar ATPase subunits responsible for acidifying endosomes as positive regulators of dextran mediated endocytosis (Supplemental tables 1 and 2). Vesicular acidification is important for inducing conformational changes in the MRC1 protein to promote ligand/dextran release (Hu *et al*., 2018).

The screen identified multiple membrane trafficking complexes that may be important for the recycling and endocytic traffic of MRC1. Following vesicular acidification, endosomes quickly merge into the conventional endo-lysosomal systems, where the CORVET and HOPS complexes mediate homotypic fusion events to consolidate vesicle cargo (Balderhaar and Ungermann, 2013). All eight members that make up the CORVET and HOPS complexes were highly ranked in the screens as positive regulators of dextran uptake, suggesting that efficient vesicle trafficking is important for endocytosis or MRC1 expression. However, many of the CORVET and HOPS subunits have additional functions regulating Golgi and autophagosome traffic, so the exact nature of their contribution is yet to be determined (Liu *et al*., 2006; Pols *et al*., 2013; Kvalvaag *et al*., 2014; Bowman *et al*., 2019; Pavlova *et al*., 2019). Of note, *Vps33a^sgRNA^* BMDM did not have a severe defect in Lucifer yellow uptake, pointing its role in MRC1 surface expression rather than homotypic fusion and endolysosomal trafficking as a possible mechanism regulating dextran uptake.

As vesicles are merged into the endolysosomal system, endocytic receptors may be recycled and returned to the surface. Rab11a-positive vesicles traffic recycled proteins to the Golgi for processing, and then Rab11a and the exocyst complex return those proteins to the surface (Takahashi *et al*., 2012). Seventy-five percent of total MRC1 protein is found in intracellular pools instead of on the surface of the cell, which suggests that receptor recycling is important for maintaining surface pools of MRC1 (Stahl *et al*., 1980; Wileman *et al*., 1984). MRC1 delivered to the surface can then be internalized again or released by extracellular proteases. The tyrosine kinase *Syk* was identified as a negative regulator of dextran uptake, which may be partly explained by its regulation of MRC1 shedding as SYK inhibition prevents MRC1 shedding by proteases and may increase MRC1 protein levels (Gazi *et al*., 2011).

In conclusion, we have developed a CRISPR/Cas9 whole genome screening method that allowed us to identify genes regulating endocytic mechanisms in primary macrophages. With this approach, we identified genes that regulate dextran uptake in BMDM. We identified MRC1-mediated endocytosis as an efficient process used by macrophages to internalize dextran at low concentrations and macropinocytosis as an efficient internalization method for higher dextran concentrations when endocytic receptors are saturated. Furthermore, we demonstrated that IL4-treated macrophages are not as highly macropinocytic as once believed. This work supports the use of Lucifer yellow as a superior marker of macropinocytosis and its use in future screens will facilitate identification of genes regulating macropinocytosis. MRC1 is a marker of alternatively activated/anti-inflammatory macrophages and is a potential target for delivery of therapeutics to macrophages (Pustylnikov *et al*., 2014). This work provides the basis for mechanistic underpinning of how MRC1 contributes to the receptor mediated uptake of carbohydrates from the tissue milieu and distinguishes genes regulating receptor mediated endocytosis from those regulating the bona fide fluid-phase uptake of fluids and solutes by macropinocytosis.

## Materials and Methods

### Reagents

Dulbecco’s modified eagle medium (DMEM) was from American Type Culture Collection (ATCC; 30-2002; Manssas, VA). Heat-inactivated fetal bovine serum (FBS) was from R&D Systems (Atlanta, GA).

Penicillin/streptomycin was from Corning (Manassas, VA). Phosphate-buffered saline (PBS), with or without calcium and magnesium, was purchased from GE Healthcare Life Sciences (Pittsburgh, PA). Primary conjugated anti-MRC1 antibody (clone C068C2), unconjugated anti-MRC1 antibody (clone C068C2), PE-conjugated isotype control IgG2a antibody (clone RTK2758), Truestain FcX (clone 93), mouse recombinant IL4, and mouse recombinant CSF1 were from BioLegend (San Diego, CA). Anti- MRC1 and anti-beta actin antibodies for immunoblots were from Cell Signaling Technology (Danvers, MA). Dylight 594 anti-rat secondary antibody (catalog No. SA5-10020) was from Thermofisher (Waltman, MA). Mannan from *S. cerevisiae* prepared by alkaline extraction was from Sigma Aldrich (St. Louis, MO). Cyclosporine A, puromycin, Lucifer yellow, Texas Red 3, 40 and 70 kDa dextran, 40 kDa amino dextran, Dylight 594 NHS ester dye, A555 NHS ester dye, A647 NHS ester dye, and NHS sulfoacetate, A647 NHS ester dye, and N,N-Dimethylformamide were from Fisher Scientific (Waltman, MA). LY294002, Dyngo-4a was from Cayman Chemical (Ann Arbor, MI). PsPAX2 was a gift from Didier Trono (Addgene plasmid # 12260 ; http://n2t.net/addgene:12260; RRID: Addgene_12260). pCMV-VSV-G was a gift from Bob Weinberg (Addgene plasmid #8454; http://n2t.net/addgene:8454; RRID:Addgene_8454). The mouse Brie CRISPR knockout pooled library was a gift from David Root and John Doench (Addgene #73633) (Doench *et al*., 2016). Sequences for sgRNAs used for targeted gene disruptions are listed in Supplemental Table S3.

### Bone marrow-derived macrophage isolation and culture

Bone marrow-derived macrophages (BMDMs) were differentiated from the bone marrow cells of C57BL/6 mice as described (Racoosin and Swanson, 1989), with some modifications (Figure 1). We used C57BL/6 mice expressing *Cas9-EGFP* from the Rosa26 locus for both whole-genome screens (Chu *et al*., 2016) (Jackson Labs Stock No. 026179, Bar Harbor, ME). For all other experiments, we used C57BL/6 mice expressing *Cas9* from the H11 locus (Chiou *et al*., 2015) (Jackson Labs Stock No. 028239, Bar Harbor, ME). Briefly, mice were euthanized with CO2. We then removed the femurs and cut off the tips to flush the bone marrow with PBS. Cells were washed and cultured on non-tissue culture treated plates in 5% CO2 and 37°C in bone marrow medium (BMM) consisting of DMEM, 20% heat-inactivated FBS, 30% L-cell Supernatant as a source CSF1(Stanley and Heard, 1977), 10,000 IU penicillin, 10 mg/ml streptomycin, and 0.0004% 2-mercaptoethanol. On day four post-isolation, non-adherent cells were discarded, and the adherent cells were cultured as macrophages in BMM. BMDMs were transduced on day 4 or 5 post isolation.

### Lentiviral transduction with Brie pooled sgRNA library

Brie library amplification, lentiviral production, and titer calculations were performed as described (Joung *et al*., 2017) with minor modifications. Briefly, the Brie library plasmids were amplified in Stbl3 *E. coli* (New England Biolabs, Ipswich, Massachusetts) and sequenced to assess sgRNA representation. To produce lentiviral particles for transduction, HEK293T cells were seeded onto 10-cm plates in 10% FBS plus DMEM before transfection with 6 µg sgRNA LentiGuide-Puro plasmid, 6 µg psPax2 plasmid, 1 µg pVSVG plasmid (Stewart *et al*., 2003), and 24 µg polyethyleneimine (PEI) for 24 h. Lentiviral supernatant was harvested after 48 h and stored at -80°C. To calculate the functional viral titer, macrophages were transduced with increasing volumes of lentivirus in BMM plus 1 µM cyclosporin A. After 48 h, the transduction solution was replaced with BMM plus 5 µg/ml puromycin for selection. After selection, the surviving cells were counted to determine the percentage of transduced cells relative to non-selected control wells. For each screen, we transduced approximately 5x10^7^ cells with a viral titer to achieve at least 50% cell death following antibiotic selection. The low titer was to ensure that the majority of transfected cells contained only one guide per cell. Cells were cultured for at least 8 days following puromycin selection to allow for gene disruption and protein turnover (Figure 1). Similar procedures were followed for lentiviral delivery of sgRNA for targeted single gene disruptions.

### Fluorescent dye conjugation to dextran

Room temperature Alexa 647, Dylight 594, or Alexa 555 NHS ester dye dissolved in DMF was added to 40 kDa amino dextran dissolved in 100 mM sodium bicarbonate buffer at a 10:1 molar ratio for at least 1 h on a rocker. Free dye was then removed using a 7000 MW cutoff Zebra Spin column (Fishersci, Waltman, Massachusetts) according to the manufacturer’s protocol. The conjugation efficiency was calculated using Beer’s law to calculate the amount of dye compared to the amount of dextran used in the initial reaction.

### Dextran uptake assays

For the whole genome screen dextran uptake assays, cells were starved overnight in CSF1-free medium (DMEM plus 10% FBS) and subsequently exposed to 300 µg/ml dextran conjugated to Dylight 594 or A555 40 kDa for 15 min in the presence (CSF1 stim), or absence (unstim) of 200 ng/ml CSF1 in DMEM plus 10% FBS preequilibrated to pH 7.4 (Figure 1). After 15 min, the cells were gently washed with warm PBS, incubated on ice for 5 min in calcium- and magnesium-free PBS, and pipetted to facilitate detachment. Detached cells were sorted with a BD FACS Jazz flow cytometer based on dextran fluorescence with the highest (high dextran uptake) and lowest (low dextran uptake) quintiles sorted for sequencing of the sgRNA inserts. Approximately 10% of the cells were collected before sorting to assess sgRNA insert distribution in the cell population (Presort). After sorting, the cells were pelleted and stored at -80°C before DNA purification. In the targeted knockout studies, dextran uptake assays were conducted with the fluorescent dextrans indicated in the figure legends and included Texas red 3, 40 and 70 kDa dextran, and A647 40 kDa dextran.

### PCR and next-generation sequencing of gRNA inserts

Genomic DNA was extracted from pellets with the GeneJET genomic DNA extraction kit (Thermofisher, Waltham, Massachusetts) and the sgRNA library was amplified for sequencing as described (Joung *et al*., 2017) (Figure 1). Briefly, genomic DNA was amplified equally in each reaction with an equimolar mix of P5 staggered primers and a unique P7 primer (Supplemental Table S4) containing an index sequence for sample identification with a Phusion polymerase kit (Fishersci, Waltman, Massachusetts). Following genomic DNA amplification, the PCR product was pooled and the 357 bp product was gel purified with the wizard gel and PCR clean up kit (Promega, Madison, Wisconsin) for sequencing with the Illumina Nextseq^TM^ 500 system high-output kit with 75 bp read length. Sequencing quality was assessed within the Illumina dashboard and read count summaries and mapping data from MAGeCK (Supplemental Figure S2 and S3). Cut adapt, version 1.18 (Martin, 2011) was used to remove sequencer adapter regions to produce FASTQ files containing the 20 bp sgRNA sequences.

### Statistical analysis of sequencing data and gene ranking

Model-based analysis of genome-wide CRISPR-Cas9 Knockouts (MAGeCK), version 0.5.9(Li *et al*., 2014) identified and ranked enriched sgRNAs in the highest and lowest quintiles based on dextran uptake. The ‘–count’ function mapped sgRNAs sequences processed with Cutadapt to the Brie library sequences file and generated read counts for each sgRNA. The ‘-test’ command was used to compare sgRNA read counts for each sgRNA in the low and high fluorescence populations and rank each sgRNA for enrichment to identify sgRNAs, and therefore genes, that regulate dextran uptake. The ‘-control-sgrna’ command was used with the ‘-test’ command to identify 1000 control non-coding sgRNAs used as a control for read count normalization.

### Fluorescence Microscopy

BMDMs cultured overnight in CSF-1-starvation medium (DMEM + 10% FBS) were stimulated with 200 ng/ml CSF1 in the presence of 100 or 500 µg/ml dextran and 500 μg/ml Lucifer yellow with the indicated inhibitors in HBSS. After each stimulation, cells were quickly washed with cold HBSS to slow biochemical activity and vesicular trafficking before imaging dextran and Lucifer yellow. Cells were pretreated with 50 µM LY294002 or 1 mg/ml mannan for 30 min before each experiment. Cells were imaged using an Olympus IX83 microscope platform at 37°C. Fluorophores were excited using the X-cite Turbo system and the emission captured by emission filters (Quad cube: OSF-QUADPLEDBX3). We observed emissions using a cooled CCD camera through a 60x oil objective (NA 1.42) or a 40x air objective (NA 0.95) (Olympus, Shinjuku City, Tokyo, Japan).

### Immunofluorescence Staining

Cells were fixed 10 days post-transduction with 4% PFA for 10 min and washed with PBS before permeabilization for 15 min with 0.3% Triton-X 100 plus 1% BSA in PBS. Then, cells were stained with anti-Mrc1 primary antibody (1:200) for 1 h in PBS plus 1% BSA. Cells were stained with anti-rat Dylight 594 secondary antibody (1:125) and DAPI for 1 h. Cells were imaged using a 40X air objective.

### Image Processing

The diameter of fluorescent dextran or Lucifer yellow puncta was measured to calculate macropinosome size. Using Fiji, the background intensity was subtracted, images were then thresholded, and the ‘Analyze particles…’ option was used to measure the Feret diameter of each punctum (Schindelin *et al*., 2012). The frequency distribution of Feret diameters was used to demarcate the size distribution of macropinosomes and plotted. The number of cells was manually tallied to calculate the number of each size puncta per cell. In each condition, >30 cells were measured.

Quantification of MRC1 from immunofluorescence image integrated intensity of anti-MRC1 stained cells were quantified using CellProfiler. The nuclei stained with DAPI and was used to segment individual cells. The bar plot shows mean integrated intensity per each genotype and the error bars are standard deviation of the mean. For each genotype >50 cells were measured. One-way ANOVA followed by Dunnett’s post hoc comparison was used to determine significance and calculate p-value.

### Surface marker staining for MRC1

A total of 1.5 x10^5^ cells were suspended in 1% FBS plus 1.5 µg/ml Fc block in PBS for 15 min on ice according to the manufacturer’s instructions. After 15 min, Phycoerythrin (PE)-conjugated anti-MRC1 or isotype control (1:80) was added to each sample and incubated for 15 min on ice. The samples were then washed three times and fluorescence intensity measured with a BD Accuri flow cytometer. At least 10^4^ cells were analyzed for each experiment and live cells gated using forward and side scatter.

### Immunoblotting

Following transduction and a culture period of ∼14 days for protein depletion, BMDMs were pelleted and lysed in ice-cold MPER (Thermo Fisher Scientific) supplemented with protease and phosphatase inhibitors, sodium orthovanadate and incubated on ice for 12 min. The lysates were spun down at 14,000 g for 15 min at 4°C. A volume of lysate yielding 20 μg total protein was loaded on to 4-20% gel and ran at 90 V for 75 min in 1% SDS in 1x TGS. Following gel electrophoresis, the protein was transferred to a PVDF blot in TGS containing 20% MeOH for 1 h at 350 mAmp on ice. Immunoblots were incubated with fluorophore-conjugated secondary antibodies recognizing the different antibody species and imaged on a Licor Odyssey Fc imaging system at 700 and 800 nm.

### IL4 treatment

Cells were stimulated with 50 ng/ml IL4 in BMM for 48 h. For experiments involving CSF1 starvation, cells were stimulated with 50 ng/ml IL4 in BMM for 24 h, and then the BMM was replaced with 50 ng/ml IL4 in 10% FBS plus DMEM for another 24 h.

### Statistical analyses

The geometric means and medians for all flow cytometry data were calculated using FlowJo software 10.6 (Franklin Lakes, New Jersey). Statistical analysis was performed using Graph Pad Prism software 8.0 (San Diego, California). We used one-way ANOVAs for the initial analysis of more than two samples followed by a Dunnet’s post hoc test for multiple comparisons.

## Conflict of Interest

The authors declare that the research was conducted in the absence of any commercial or financial relationships that could be construed as a potential conflict of interest.

## Author Contributions

JW, NT and AH conceived of the project. JW, KW, LH, BP and NT planned the experiments. JW, JK, LH, KW, BP and NT executed the experiments. JW, LH, KW, AH and NT analyzed the data and generated the figures. JW generated the first draft of the manuscript. JW, NT and AH wrote and edited sections of the manuscript. All authors contributed to manuscript revision and approved the submitted version.

## Funding

Research reported in this publication was supported by the National Institute Of General Medical Sciences of the National Institutes of Health under Award Number R15GM139162 and Award Number P20GM135008. The content is solely the responsibility of the authors and does not necessarily represent the official views of the National Institutes of Health. This material is based upon work conducted using the SDSU Genomics Sequencing Facility (RRID:SCR_023959) and the SDSU Functional Genomics Core Facility (RRID:SCR_023786) supported in part by the National Science Foundation/EPSCoR Grant No. 0091948 and No.IIA-1355423, the South Dakota Agricultural Experiment Station, and by the State of South Dakota. Any opinions, findings, and conclusions or recommendations expressed in this material are those of the authors and do not necessarily reflect the views of the National Science Foundation (USA) or National Institutes of Health (USA).

### Acknowledgements

The authors thank Joel Swanson for his suggestions and edits to the manuscript.

## Data Availability Statement

The datasets generated for this study can be found in the GEO repository, accession number GSM7623687.

## Abbreviations

BMDM: bone marrow-derived macrophages
*Mrc1*/MRC1: mannose receptor c-type 1
CSF1: colony stimulating factor-1
CSF1R: colony stimulating factor-1 receptor
FDR: false discovery rate
LFC: log2-fold change

---

Supplemental Table S1. MAGeCK low/high test results from the unstimulated dextran uptake CRISPR/Cas9 whole genome screen

*See attached excel file*.

Supplemental Table S2. MAGeCK low/high test results from the CSF1-stimulated dextran uptake CRISPR/Cas9 whole genome screen

*See attached excel file*.

**Supplemental Table S3.**
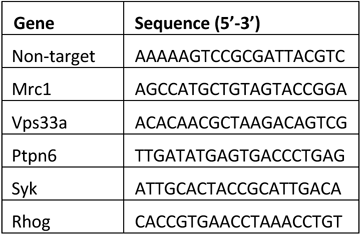
sgRNA sequences used for targeted gene disruption.

**Supplemental Table S4.**
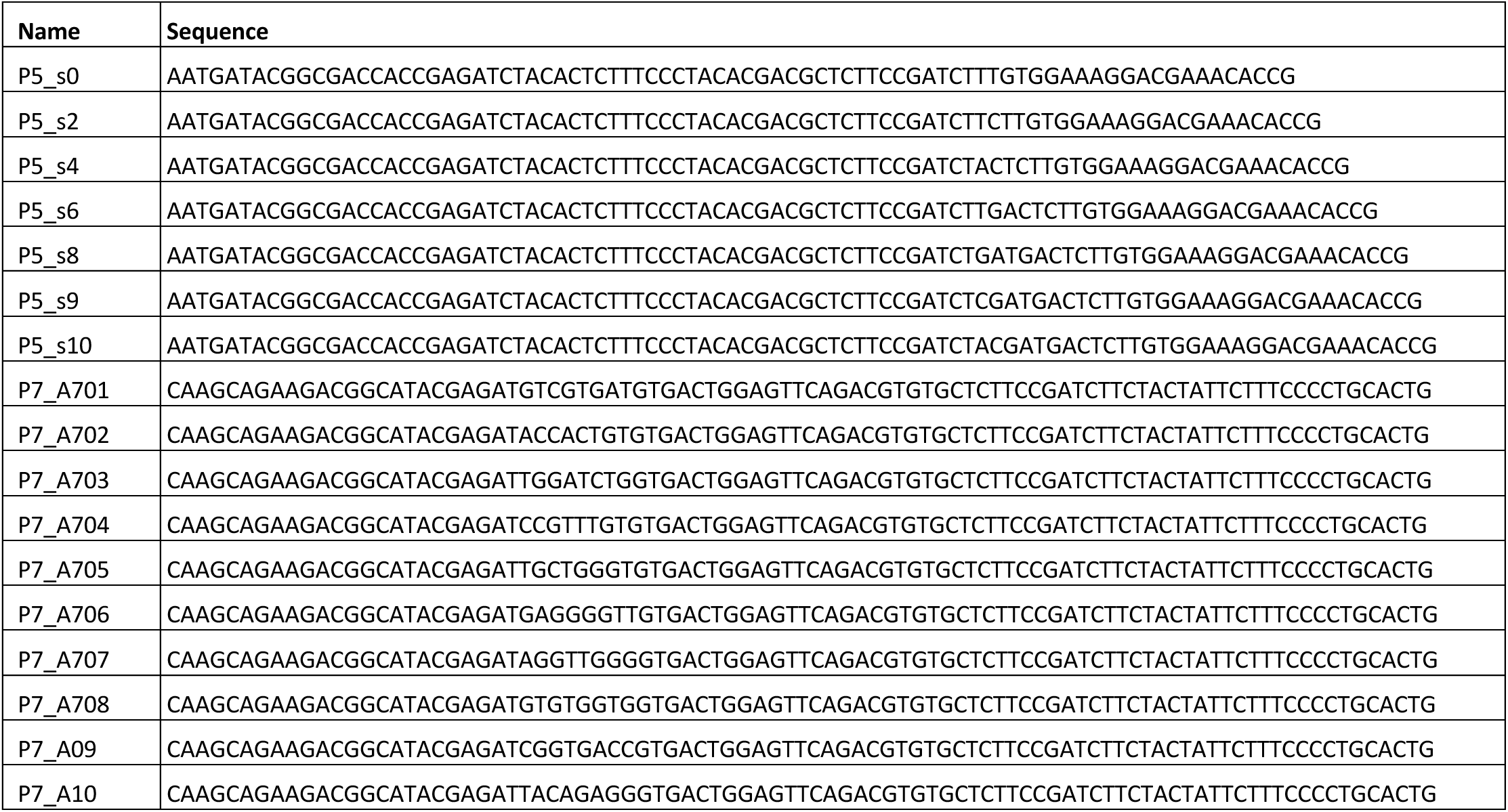
Primers sequences used to amplify sgRNA inserts from genomic DNA for next generation sequencing.

**Supplemental Figure S1.**
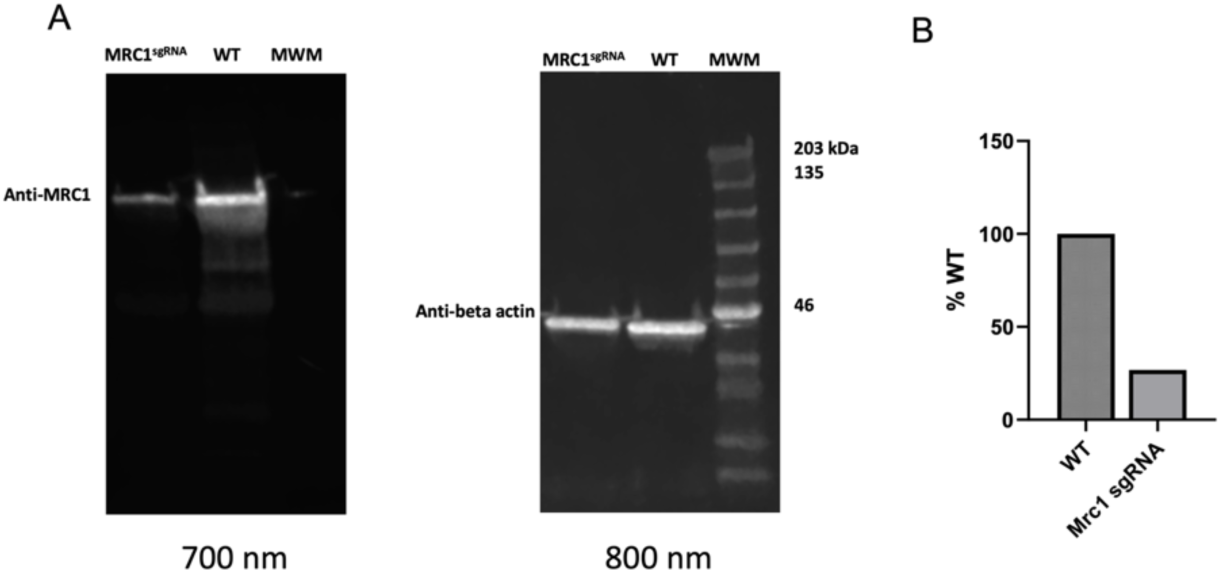
Western blot of Mrc1^sgRNA^ transduced BMDM. BMDM transduced with Mrc1 sgRNA and WT BMDM were lysed and run on SDS-PAGE gel followed by transfer to PVDF membrane. Immunoblots were incubated with secondary antibodies recognizing the different hosts of the anti- MRC1 and anti-actin antibodies. B. Intensities of the bands were quantified in ImageJ, and percent WT values were plotted. MWM = molecular weight marker

**Supplemental Figure S2.**
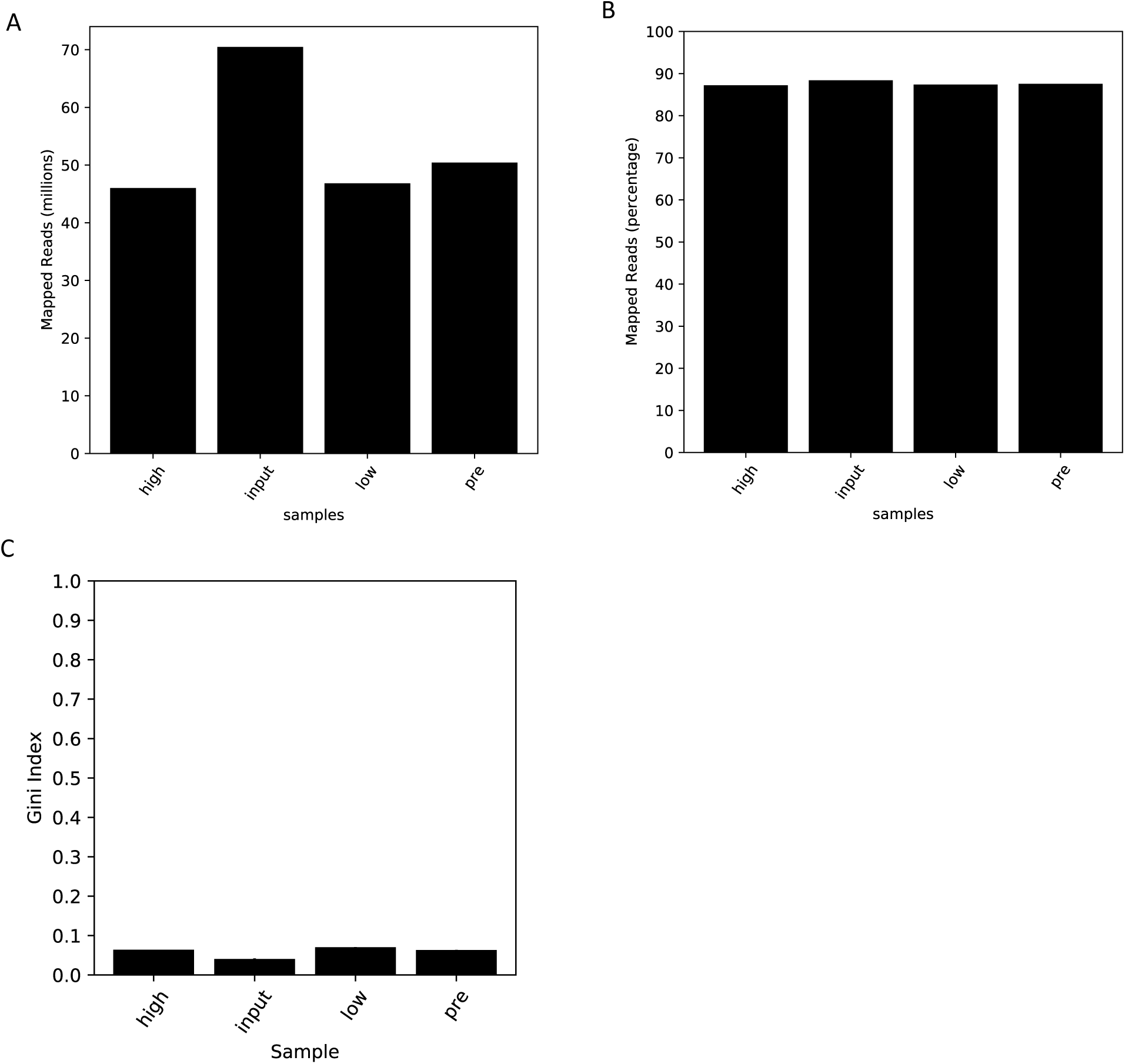
Quality control data describing MAGeCK read mapping for CSF1 stimulated dextran uptake screen. A) This bar graph shows the number of mapped reads for each sample. B)This bar graph describes what percent of reads were mapped within each sample. C) This bar graph describes the Gini Index for each sample. The Gini index is a measure of inequality. 1.0 indicates 1 sgRNA has all of the reads while 0.0 would indicate the reads are equally distributed amongst all sgRNAs.

**Supplemental Figure S3.**
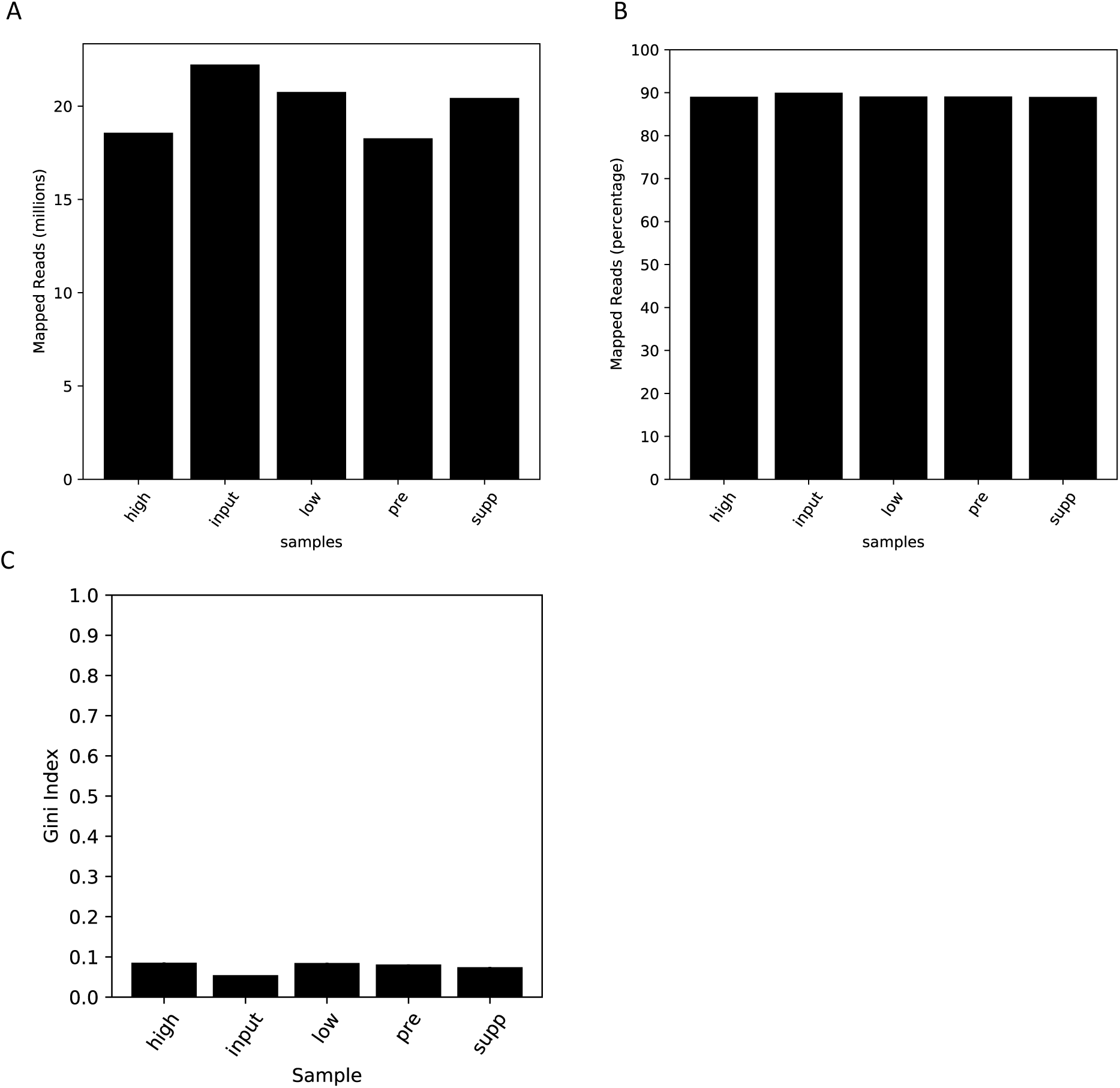
Quality control data describing MAGeCK read mapping for unstimulated dextran uptake screen. A) This bar graph shows the number of mapped reads for each sample. B) This bar graph describes what percent of reads were mapped within each sample. C) This bar graph describes the Gini Index for each sample. The Gini index is a measure of inequality. 1.0 indicates 1 sgRNA has all of the reads while 0.0 would indicate the reads are equally distributed amongst all sgRNAs.

## Notes

https://www.ncbi.nlm.nih.gov/geo/query/acc.cgi?acc=GSM7623687

### Competing Interest Statement

The authors have declared no competing interest.

